# Characterizing intra-tumor and inter-tumor variability of immune cell infiltrates in murine syngeneic tumors

**DOI:** 10.1101/2020.12.22.423997

**Authors:** Sepideh Mojtahedzadeh, Alan Opsahl, Joan-Kristel Aguilar, Dingzhou Li, Nicole Streiner, Jinwei Wang, Dusko Trajkovic, Germaine Boucher, Timothy Coskran, Shawn P. O’Neil, Sripad Ram

**Author notes:** These authors contributed equally (Sepideh Mojtahedzadeh, Alan Opsahl).

## Abstract

By using serial-sectioning methodology combined with immunohistochemistry (IHC) and whole-slide digital image analysis (DIA), we present a systematic analysis of intra-tumor and inter-tumor variability in the abundance of nine immune cell biomarkers in multiple murine tumor models. Our analysis shows that inter-tumor variability is typically the dominant source of variation in measurements of immune cell densities. Statistical power analysis reveals how group size and variance in immune cell density estimates affect the predictive power for detecting a statistically meaningful fold-change in immune cell density. Inter-tumor variability in the ratio of immune cell densities show distinct patterns in select tumor models and reveal the existence of strong correlations between select biomarker pairs. Further, we show that the relative proportion of immune cells at different depths across tumor samples is preserved in some but not all tumor models thereby revealing the existence of compositional heterogeneity. The results and analyses presented here reveal the complex nature of immunologic heterogeneity that exists in murine tumor models and provide guidelines for designing preclinical studies for immuno-oncology research and drug development.

## INTRODUCTION

The advent of immunotherapy has generated significant interest in characterizing immune cell infiltrates in the tumor microenvironment. A hallmark of most cancer indications is the presence of intratumor heterogeneity which is recognized as a key factor contributing to poor prognosis, treatment failure and drug resistance [1, 2]. Tumor heterogeneity has traditionally been characterized in terms of tumor-intrinsic genetic variations at the clonal and sub-clonal levels [3, 4]. Recently, immunologic heterogeneity within the tumor microenvironment has received a lot of attention as it has been shown to be a major predictor of phenotypic heterogeneity and treatment response to immunotherapeutic agents [5]. Murine syngeneic tumors represent the most widely used model system for preclinical drug discovery and development [6, 7]. Several studies have characterized murine syngeneic tumors, which have revealed various facets of the tumor-immune contexture such as the genomic and transcriptomic landscape, mutational burden and immunogenicity, and the composition of different immune cell subtypes [8–11].

Intratumor heterogeneity is typically assessed through regional measurements in bulk tissue (e.g biopsy of primary tumor versus metastatic lesion) using grind and bind approaches (e.g. RNASeq, flow cytometry, etc.), and consequently lack spatial information about the cells [9, 10, 12]. Intratumor heterogeneity has also been investigated in thin (4-20 microns) tumor sections which facilitates assessment in pathologically distinct regions within the tumor section [8, 11]. However, as tumor tissue is a three-dimensional structure, an important question arises as to whether the measurement of immune cell abundance from a thin tumor section is preserved across different depths within the tumor. Specifically, the depth dependent variation in immune cell density and/or its composition within murine tumors has received very little attention.

Here, we present a comprehensive analysis to quantify the heterogeneity in the density and composition of nine immune cell biomarkers (CD3, CD4, CD8α, CD11b, CD45, F4/80, FoxP3, GR1 and Granzyme B) at different depths within multiple murine tumor models. We consider 4 different models, including ectopic allografts of CT26 and EMT6 tumors implanted subcutaneously, an orthotopic allograft of a murine pancreatic tumor (KPC-ortho) and a genetically engineered murine (GEM) tumor model of pancreatic cancer (KPC-GEM). CT26 and EMT6 are cell-line derived tumor models that have synchronous tumor growth and are easy to handle, which make them appealing as preclinical model systems. However, these models are non-autochthonous and their rapid growth rates make them poor models of human tumors. On the other hand, the KPC-GEM is autochthonous and thus more translationally relevant to human tumors. However, the asynchronous growth and labor-intensive management of these tumors make this a challenging model for routine and repeated use, such as efficacy studies for drug development. The KPC orthotopic allograft is a cell-derived model from the KPC-GEM model that is implanted at the site of origin (i.e. the pancreas) and thus represents an intermediate model among those considered here in terms of translational relevance.

We characterize heterogeneity by calculating the intra-tumor variability and inter-tumor variability for all the immune cell biomarkers. The former pertains to variability in immune cell abundance at different depths within a tumor tissue, while the latter pertains to variability in immune cell abundance among tumor specimens within a given tumor model. Our analysis reveals that inter-tumor variability is typically the dominant source of variation in all the tumor models tested here. We present a statistical power analysis that is based on retrospective IHC-DIA data, which reveals the minimum sample size (i.e. number of animals / treatment group) that is required to detect a statistically meaningful fold-change in cell density for the various immune cell biomarkers considered here.

We also investigate inter-tumor and intra-tumor variability in the ratio of immune cell densities which show distinct behavior from that of the individual immune cell densities. Specifically, our analysis reveals strong correlation in immune cell abundance between T-cell related biomarkers in select tumor models, which in turn affect the variability in the corresponding immune cell ratios. Further, we investigate the relative proportion of the various immune cell biomarkers considered here and show that the immune cell composition is well maintained among tumors in some but not all tumor models.

The results and analysis presented here provide novel insights into the nature and extent of immunologic heterogeneity in murine tumor models. Further, they reveal model-specific differences in the density and relative proportion of immune cells. We anticipate that these results will provide guidelines in designing invivo studies using murine tumor models for drug discovery and development efforts.

## METHODS

### Animal models

All animal procedures complied with the Guide for the Care and Use of Laboratory Animals (Institute for Laboratory Animal Research, 2011) and were approved by the Pfizer Global Research and Development Institutional Animal Care and Use Committee (IACUC). The KPC (KRas^G12D^/Trp53^R172H^/Pdx1-Cre) genetically engineered mouse model of pancreatic cancer [13] was generated in-house. Briefly, Trp53^LSL-R172H/+^(Jax 008652), Kras^LSL-G12D/+^(Jax 008179) and Pdx-1-Cre (Jax 01467) strains were interbred to obtain the triple mutant animals which had a strain background that was 85% B6.FVB and 15% 129/SvJae. The animals were maintained in a pathogen-free facility and routinely palpated to detect the presence of tumors. Approximately 7 to 10 days after detecting a palpable tumor the animals were euthanized and the tumors were harvested.

A KPC tumor cell line was generated as described previously [14], but with some modifications. Briefly, tumor tissues explanted from KPC mice were washed in PBS with 20 μg/ml Gentamycin, minced with a sterile scalpel in a petri-dish containing complete medium (DMEM/10%FBS/2mM L-Glutamine/1x NEAA/1x Sodium Bicarbonate/10mM HEPES/ 1x Pen/Strep) and then incubated with complete medium containing 1 mg/ml Collagenase P (Roche) for 90 minutes at 37°C. The tumor tissue was then washed twice in complete media, filtered through a 40 micron strainer to create a single cell suspension, which was cultured in complete medium containing 400 ng/ml cholera toxin for the first 7 passages. After the tenth passage, the cells were switched to normal complete medium DMEM plus 10% FBS plus 1x Pen/Strep and maintained at 37°C in a humidified incubator.

For orthotopic implantation of KPC cells, a left subcostal incision into the peritoneal cavity was made and the pancreas was externalized. One million KPC tumor cells in 50 μl PBS was slowly injected into the head of the pancreas with 27 gauge or smaller needle. The incision was closed in two layers. First, the muscle layer was apposed with 6-0 surgical nylon suture, and then the skin layer was closed with 5-0 suture or surgical wound clips. CT26 and EMT6 cell lines were purchased from ATCC. Female BALB/c mice were purchased from Jackson laboratories (Sacramento, CA). Cell-line authentication and mycoplasma testing was periodically carried out for all cell lines. CT26 or EMT6 cells were subcutaneously implanted into the dorsal side of BALB/c mice. Tumor size was measured using an electronic caliper three times per week and the tumor volume was determined using the formula: (length × width^2^)/2. Once tumor size reached ~400-500 mm^3^, the mice were euthanized, and the tumors were harvested.

All tumor samples were fixed in 10% neutral buffered formalin for 48 hours at room temperature. The tumors were then trimmed such that a central region with maximal cross-sectional area was exposed which was then embedded *en face* in paraffin blocks.

### Immunohistochemistry

Five micron sections of tumors were immunolabeled with one of the following antibodies using a Leica Bond III automated immunohistochemistry instrument (Leica Biosystems, Buffalo Grove, IL): Rabbit anti-CD3, 1:1000 or 0.2μg/ml (ab5690, Abcam, Cambridge, MA); Rat anti-CD4, clone 4SM95, 1:400 or 1.25μg/ml (14-9766-82, Thermo Fisher Scientific, Waltham, MA); Rat anti-CD8α, clone 4SM15, 1:2000 or 0.25μg/ml (14-0808-82, Thermo Fisher Scientific); Rabbit anti-CD11b, clone EPR1344, 1:22000 or 0.058μg/ml (ab133357, Abcam); Rabbit anti-CD45, 1:2500 or 0.36μg/ml (ab10538, Abcam); Rat anti-F4/80, clone BM8, 1:15000 or 0.033μg/ml (13-4881-82, Thermo Fisher Scientific); Rat anti-FoxP3, clone FJK-16s, 1:200 or 2.5ug/ml (14-5773-82, Thermo Fisher Scientific); Rat anti-GR-1, clone RB6-8C5, 1:9000 or 0.055μg/ml (14-5931-82, Thermo Fisher Scientific); Rat anti-Granzyme B, clone 16G6, 1:6000 or 0.083μg/ml (14-8822-82, Thermo Fisher Scientific).

Briefly, tissue sections were loaded onto the Leica Bond III instrument and deparaffinized, followed by pretreatment with Epitope Retrieval Solution 2 (AR9649, Leica Biosystems) for 20 minutes and then blocking buffer for 10-20 minutes. All rat primary antibodies were incubated for either 20 or 30 minutes followed by biotinylated rabbit anti-rat linker antibody (BA-4001, Vector Labs, Burlingame, CA) applied at either 1:300 or 1:500 for 15-30 minutes prior to detection and color visualization using Refine DAB Polymer (DS9800, Leica Biosystems). Rabbit primary antibodies were incubated for either 15 or 30 minutes followed by detection with Refine DAB Polymer. All slides were counter stained with hematoxylin included in the Refine DAB Polymer kit.

To minimize operator variability, specific individuals were assigned to section the paraffin blocks and execute all IHC assays for a given tumor model. To ensure consistency, all IHC assays were conducted on the same automated immunolabeling instrument and all slides were scanned using the same whole slide digital scanner. For the precision study, 15 adjacent serial sections were prepared for each biomarker from a CT26 tumor block; consequently, these sections were immunolabeled, scanned and analyzed in the same manner as the study samples.

### Slide scanning and digital image analysis

All slides were scanned using an Aperio AT2 whole-slide digital scanner (Leica Biosystems, Vista, CA) at 20x magnification. Whole-slide, automated image analysis was performed using Visiopharm software (Visiopharm, Hoersholm, Denmark). The two DIA endpoints, 1) immune cell density and 2) IHC area%, were calculated separately by different personnel in a semi-blinded manner. Immune cell density is a cell-based metric that relies on the detection of individual cells that are positive for the biomarker and is defined as the ratio of the number of marker-positive cells to the viable tumor area (expressed in 1/mm^2^). IHC area% is a pixel-based metric that relies on the total area of pixels in the IHC image that are positive for the biomarker, defined as 100 times the ratio of the total area of the biomarker-positive pixels to the viable tumor area.

For each tumor model, custom Visiopharm APPs were developed to detect the tissue in the image, to delineate viable tumor and non-tumor regions, and to compute the viable tumor area. In addition, areas of necrosis and hemorrhage, identified on H&E images from serial sections, were manually excluded from subsequent analysis. For each biomarker two custom apps were implemented, i.e., the cell count app and the IHC area% app. The cell count app detects and counts cells that are positive for the biomarker of interest in the viable tumor regions. The IHC area% app detects all pixels that are positive for the chromogen (DAB) used to localize the biomarker of interest in the viable tumor regions and then outputs the total area of the positive pixels.

### Data analysis

The calculation of inter-tumor and intra-tumor coefficient of variation (CV) was carried out using MATLAB (Mathworks, Natick, MA). For each biomarker, we calculated a robust estimate of the CV which is given by [15]:

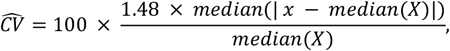

where X = (x_1_,x_2_,…x_N_) denotes the data and x ∈ *X*. We note that the above definition of CV is different from the conventional definition of CV which is given by the ratio of the standard deviation to the mean multiplied by 100. A shortcoming of conventional CV is that its numerical value increases when the measurements (x_1_, x_2_, …, x_N_) are relatively small values (i.e. ≤ 1). In many cases, a lower limit of quantification is empirically set and measurement values below this limit are censored from the CV calculation which results in a biased CV estimate [16]. By using a robust estimate of CV as defined above we avoid censoring the data and provide a more accurate estimate of CV.

Heatmaps of immune cell density estimates (Figure 2), immune cell ratios (Figure 4) and the relative proportion of immune cells within and among tumor samples (Figure 5) were generated using MATLAB. The calculation of Spearman’s correlation coefficient between cell density estimate and IHC area% (Supplementary Figures 4-7) and linear regression analysis (Supplementary Figure 8) were carried out using Prism v8.4.2 (GraphPad, San Diego, CA). Spearman’s correlation coefficient between different pairs of biomarkers (Table 1 and Supplementary Table 1) was calculated using MATLAB. These results were then exported to Excel and displayed as an upper triangular matrix where the values were color coded to indicate the degree of correlation.

**Table 1.**
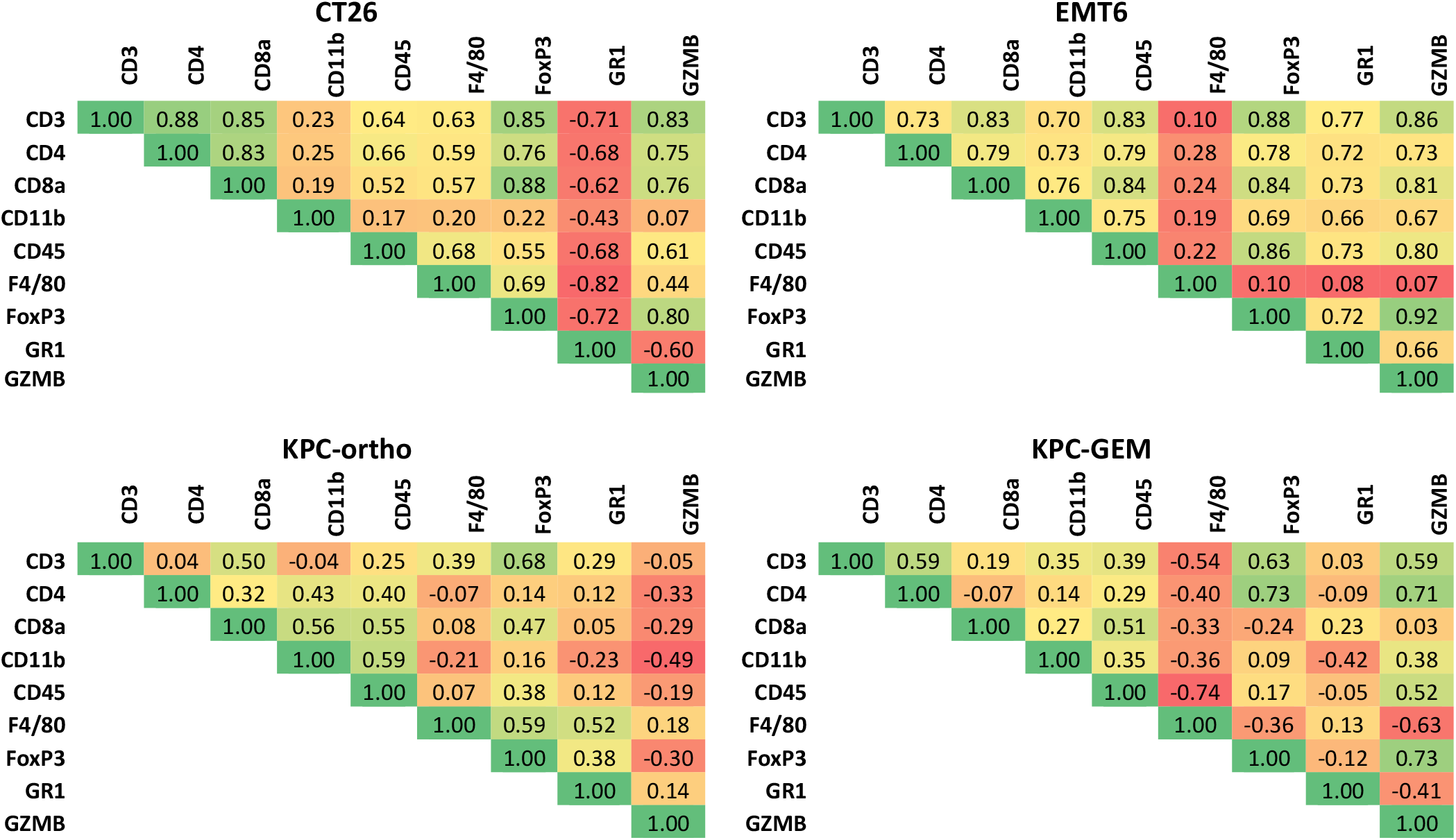
The table lists the Spearman’s correlation coefficient between immune cell density estimates for a pair of biomarkers in different tumor models. The color coding from green to red indicates high positive correlation (+1) to high negative correlation (−1).

### Retrospective analysis of IHC-DIA data

We retrieved data pertaining to the vehicle-control group from 12 different preclinical studies involving murine syngeneic tumors (CT26, MC38 and KPC-GEM tumors; n = 5 – 10 animals per group) that were performed in-house between January 2016 and November 2017. Specifically, these studies were selected since the same IHC assays and whole-slide image analysis workflows described here were used with these study samples, and IHC-DIA was performed on a single section from every tumor. The CV for each biomarker was calculated using the above formula by pooling the cell density estimates for that biomarker from all 12 studies.

### Statistical power analysis

Power analysis was conducted using a two-sample, non-central t-test paradigm with standard deviations estimated from the retrospective. A range of sample size, desired power, and detectable effect size were used in the calculation. The significance level was set to 5% (i.e., p<0.05).

## RESULTS

We extensively use IHC combined with whole-slide DIA analysis in our internal programs to assess biomarker abundance in murine syngeneic tumors. The motivation to investigate tumor heterogeneity arose due to a retrospective analysis of our internal data where we observed high variability in immune cell density estimates within the mock/vehicle-control treated groups across multiple studies (Figure 1A). Specifically, we observed that the estimate of CV for most immune cell biomarkers was above 20%, a cutoff value that is commonly used to assess variability in IHC assays [17–19]. We hypothesized that the high CV in immune cell density estimates could be attributed to two sources, i.e. inter-tumor and intratumor variability, which likely arise due to the stochastic nature of biological processes that exist even in genetically identical, inbred mouse strains. As IHC methodology is intrinsically a two-dimensional technique in that it samples a very thin (4 – 20 microns) region from a thick tumor tissue, a question arises as to whether inter-or intra-tumor variance is the dominant source of variability that is contributing to the high CV of immune cell density estimates.

**FIGURE 1.**
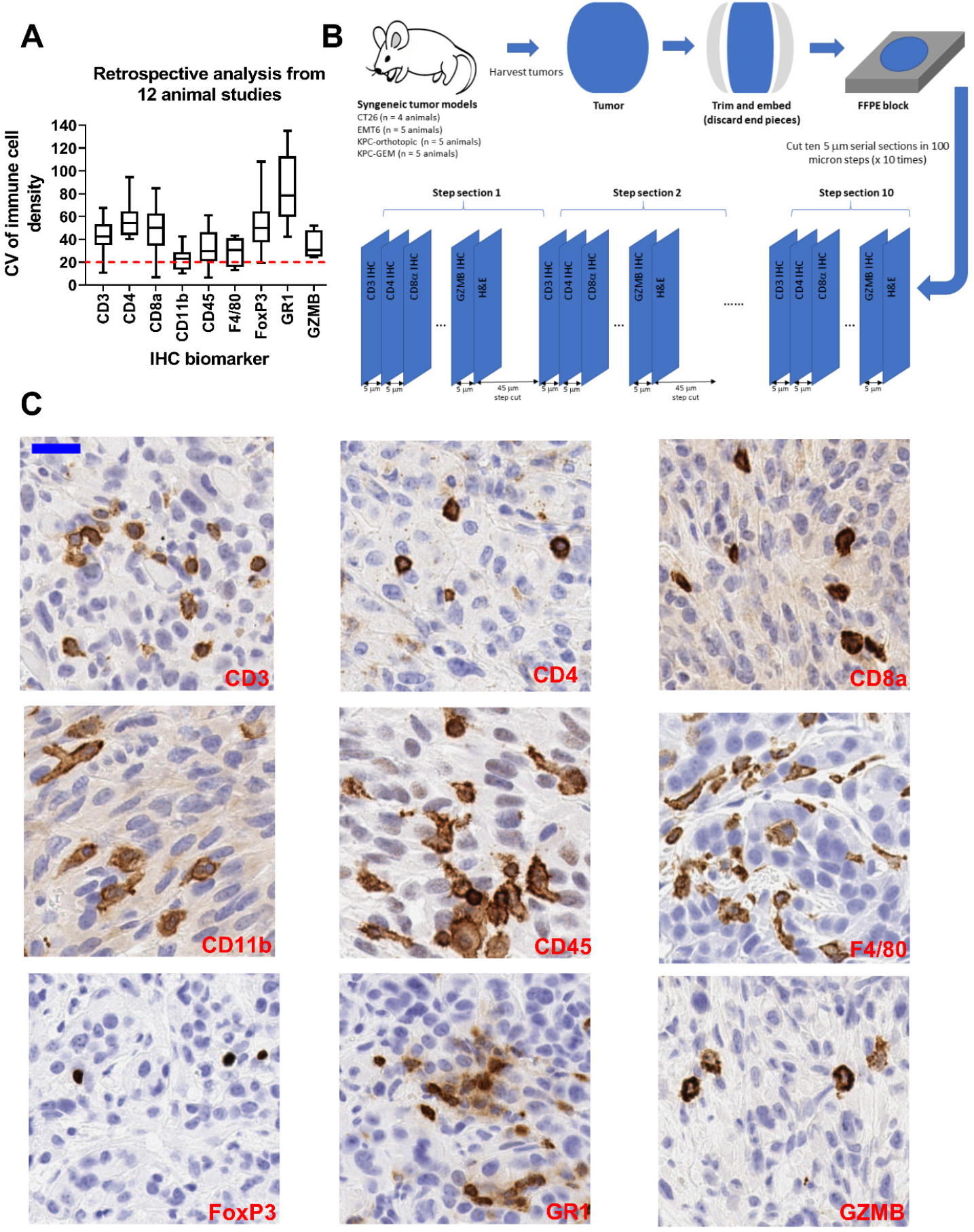
Retrospective analysis of animal-to-animal variability in immune cell density estimates. Panel A shows the coefficient of variation (CV) of immune cell density estimates for nine biomarkers quantified through IHC and whole-slide digital image analysis. The CV for each biomarker was calculated using data from vehicle control groups taken from twelve different animal studies involving murine syngeneic tumor models. Panel B shows the study design to quantitatively assess intra-tumor and inter-tumor variability in four different murine tumor models. Tumors were grown in mice, harvested, trimmed and archived as formalin-fixed, paraffin-embedded (FFPE) blocks. For each tumor sample, a set of ten serial sections (each 5 microns thick) were cut at 100-micron step intervals up to a depth of 1 mm. Each serial section at a given step was assigned to a biomarker and the order of assignment was maintained in all the steps. In this way, the abundance of each biomarker in the tumor tissue was sampled across ten sections that were each 100-microns apart and had similar tissue cross-sectional area. Panel C shows representative regions cropped from IHC images of the immune cell biomarkers considered in this study. The biomarkers of interest were immunolabeled and detected with the brown chromogen diaminobenzidine (DAB). Tissue sections were counter-stained with hematoxylin (blue) to label nuclei. All images were cropped from whole-slide scans at 20x magnification. Scale bar equals 25 microns.

### Experimental strategy to assess intra-tumor and inter-tumor variability

To investigate the different sources of variability, we designed an experimental strategy to interrogate biomarker expression at different depths in tumors (Figure 1B). Specifically, tumor tissue was sampled at 100 micron steps up to a depth of 1 mm, and at each step at least ten 5-micron serial sections were cut. Intra-tumor variability was assessed by performing IHC followed by whole-slide DIA analysis for each immune cell biomarker on a set of 10 sections, each 100 microns apart, from a tumor sample (Figure 1B). Specifically, for every biomarker we calculate the CV of the DIA endpoint (cell density or IHC area%) from the 10-step sections obtained from each tumor sample. Intra-tumor variability was then defined as the median of the CV of the DIA endpoint from different tumor samples for a given tumor model. Inter-tumor variability was assessed by comparing the IHC-DIA results among different tumor samples within a given model. Specifically, inter-tumor variability was defined as the CV of the DIA endpoint from all step sections among all tumor samples within a given model.

To rule out IHC assay reproducibility as a potential source of variability, we performed a precision study for which 15 serial sections from a CT26 tumor specimen were subjected to our IHC-DIA workflow for each biomarker target. IHC assay-related variability for each immune cell biomarker was then assessed by calculating the CV of DIA endpoints measured from these serial sections which was found to be less than 20% for most of the immune cell biomarkers (Supplementary Figure 1).

### Intra-tumor and inter-tumor variability of immune-cell biomarkers

Figure 2A shows the heatmap of immune cell density estimates for the nine biomarker targets in the EMT6 tumor model. For display purposes, the cell density estimate for each biomarker is normalized to one so that the data for all the biomarkers can be displayed in the same heatmap. The variation in cell density estimates for each biomarker was relatively low within a given tumor, with intra-tumor variability values of less than 25% for most of the biomarkers evaluated (Figure 2C). An immediate implication of this observation is that the density of these immune cell biomarkers is relatively consistent across different depths within the sampled tumor tissue. In contrast, there is considerable variation in the cell density estimates for each biomarker among different tumors (Figure 2A and 2B) in the EMT6 tumor model. For example, the average cell density for CD3 in tumors 4 and 5 is two to four-fold lower than the average cell density for tumors 1 – 3 (Figure 2B). Consequently, the inter-tumor variability for all biomarkers was consistently higher than the corresponding intra-tumor variability, with CV values exceeding 50% for most biomarkers. This suggests that animal-to-animal variability is a major contributor to the variation in cell density estimates across different tumors.

**Figure 2.**
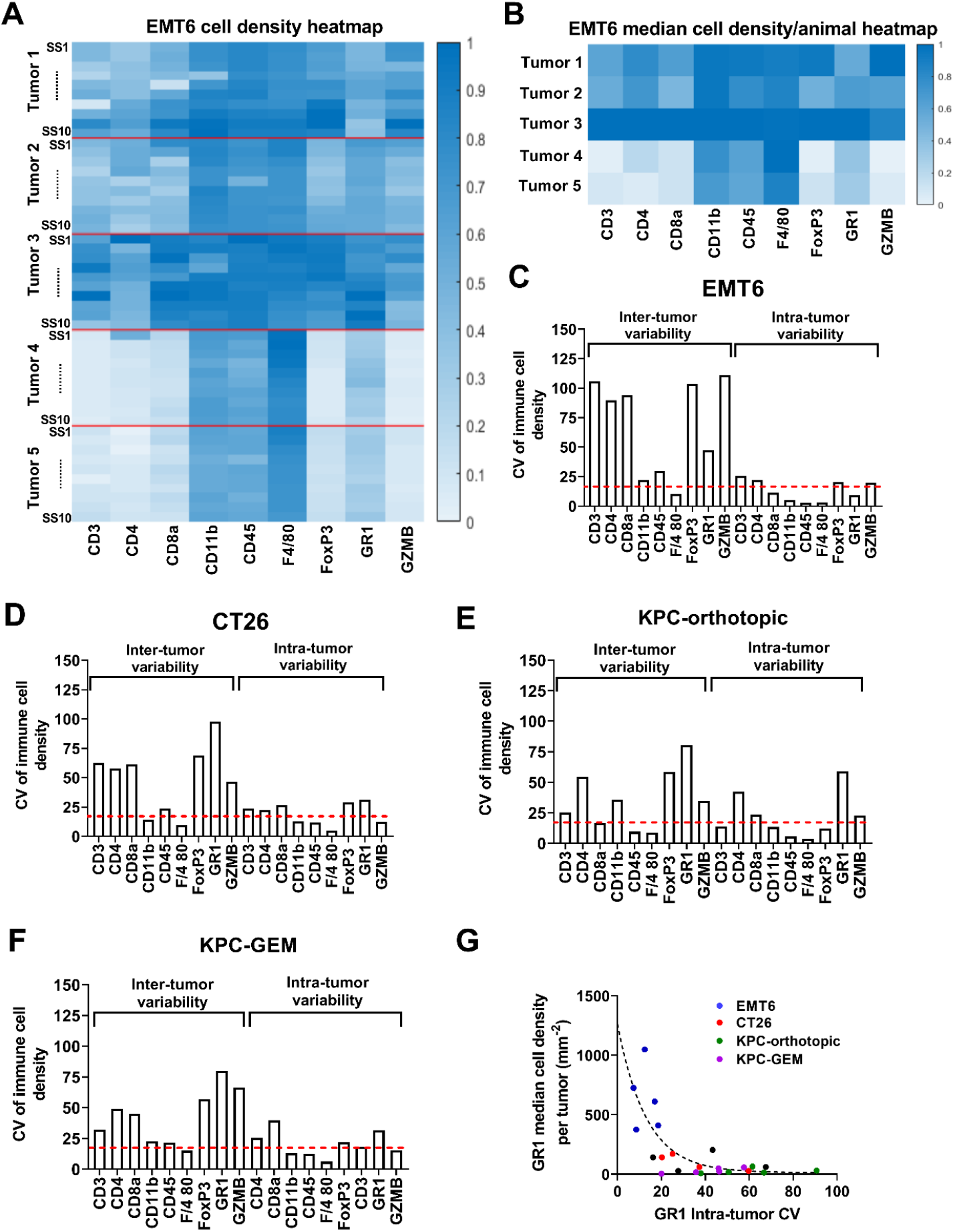
Intra-tumor and inter-tumor variability of immune cell abundance in syngeneic murine tumor models. Panel A shows the heatmap of immune cell density estimates pertaining to the EMT6 tumor model for nine biomarkers across all step sections (SS) and animals. For display purposes, the immune cell density for each biomarker is normalized to 1 for all animals; thus, the results for all biomarkers can be compared using a global colormap. The heatmap color coding is comparable within tumor specimens for a given biomarker but not between biomarkers. Panel B shows the normalized heatmap of the median immune cell density for each biomarker in the EMT6 tumor model, calculated as the median of the ten step sections obtained from each tumor. The heatmap is normalized in a manner that is analogous to the heatmap shown in Panel A. Panels C, D, E and F show the quantification of intra-tumor and intertumor variability for EMT6, CT26, KPC-orthotopic and KPC-GEM murine tumor models, respectively, for the nine immune cell biomarkers evaluated in this study. Panel G shows the plot of median cell density per animal versus intra-tumor CV for GR1 among different tumor models. An inverse relationship was observed between median cell density and the associated CV. Tumors with high cell density estimates tended to have lower intra-tumor CV, while tumors with low cell density estimates had higher intra-tumor CV.

A similar pattern of intra-tumor versus inter-tumor variability was also observed for each biomarker in the other tumor models (Figures 2D – 2F). A notable exception was found for GR1, for which the intra-tumor variability was greater than 20% in all tumor models except EMT6. This result can be attributed to the inverse relationship between intra-tumor CV and the density of GR1+ cells in the tumor tissue (Figure 2G). Specifically, the median GR1 cell density per tumor for the EMT6 tumor model is consistently higher than that for the other tumor models. In contrast, for CD45, CD11b and F4/80, intra-tumor and inter-tumor variability are consistently low in all the tumor models. This suggests that these biomarkers are uniformly expressed throughout the tumor specimen in each of the tumor models, and consequently the variability within and among tumor samples is consistently low for a given tumor model. In general, we observed that biomarkers with low intra-tumor (or low inter-tumor) CV have relatively high immune cell density estimates, resulting in an inverse correlation between median cell density per tumor and intra-tumor CV (Supplementary Figure 2).

Similar patterns were observed for CV estimates of IHC area% in that inter-tumor variability was consistently higher than intra-tumor variability for all biomarkers for each of the tumor models evaluated (Supplementary Figure 3). Furthermore, we observed a statistically significant positive correlation between cell density and IHC area% for all biomarkers in all tumor models evaluated (Supplementary Figures 4 – 7).

### Statistical power analysis reveals optimal group size for in vivo studies

Our analysis shows that inter-tumor variability is typically the dominant source of variation in cell density estimates for the immune cell biomarkers we evaluated. Given this finding, we sought to estimate the optimal group size (i.e. number of animals/group) to enable quantification of IHC-DIA endpoints capable of detecting statistically significant fold-changes in immune cell densities among groups in studies using murine syngeneic tumors. Using the CV estimates from our retrospective IHC-DIA data (Figure 1A), we carried out a power analysis to understand the relationship between detectable fold change and sample size at different power levels. A power analysis provides a measure of the likelihood that a study will detect an effect (for example, increased immune cell infiltration) if it is present.

Figure 3 shows the power analysis results for the nine immune cell biomarkers. evaluated. As expected, there was an inverse dependence between sample size and statistically meaningful, fold-change in cell density that could be detected for each biomarker. For instance, for CD3, our analysis predicts that a sample size of 4 animals will enable us to detect a 3-fold or higher change in cell density estimate at 80% power. In contrast, if we increase the sample size to 8 animals then we will be able to detect a 2-fold or higher change in cell density estimate at the same power level. Note that the minimum sample size to detect a certain fold-change in cell density varies for each biomarker. This can be attributed to the different CV values associated with each biomarker. For instance, for CD11b (which has a median intertumor CV of ~20% from our retrospective data), the power analysis predicts that a sample size of 4 animals is sufficient to detect a 2-fold or higher change in cell density at 80% power. In contrast, for FoxP3, which has a median inter-tumor CV of ~45%, our analysis predicts that a sample size of at least 10 animals per group is required to detect a 2-fold or higher change in cell density.

**Figure 3.**
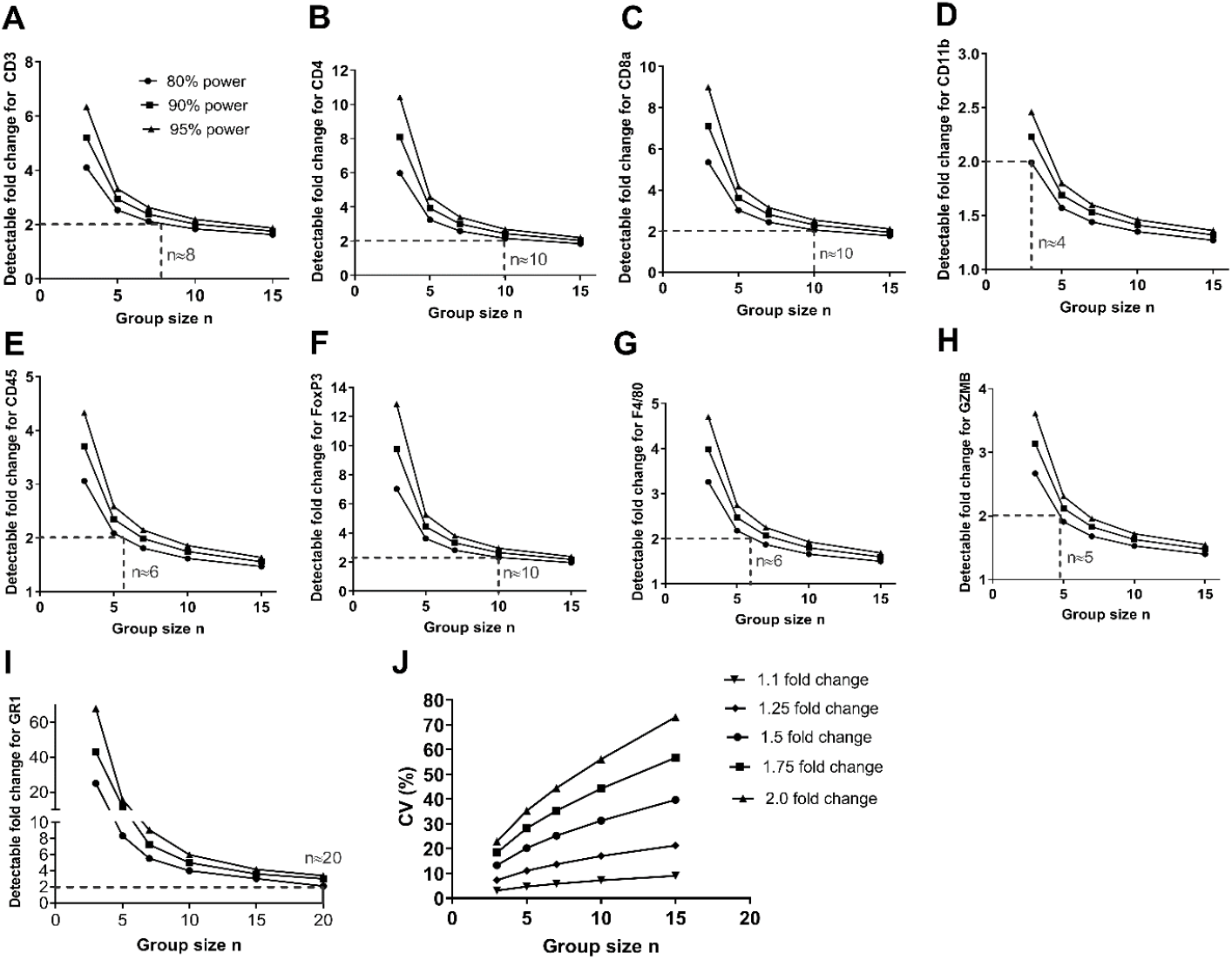
Results of statistical power analysis for IHC-DIA cell density estimates. Panels A though I show the results of power analysis for different immune cell biomarkers to quantify the dependence of group size on statistically meaningful, detectable fold change in immune cell density estimates. For the power analysis calculations, the CV for each biomarker from the retrospective analysis (shown in Figure 1A) was considered at statistical power levels of 80%, 90% and 95%. The number of animals required to detect a 2-fold difference in cell density estimates at 80% power is indicated for each biomarker. Panel J shows the relationship between CV and group size for different fold-changes in cell density, estimated at the 80% power level.

We next investigated the relationship between CV and sample size for different fold-change levels that one may seek to detect in a study for any given immune cell biomarker (Figure 3J). There is a direct relationship between the desired sample size and CV of biomarker to detect a certain fold change in cell density. For instance, we require a minimum of 10 animals/group to detect a 1.5-fold change in cell density with statistical significance when the biomarker CV is 30%. If the biomarker CV is 15 %, then we will require ~7 animals/group to detect the same fold-change in cell density with statistical significance.

### Intra-tumor and inter-tumor variability in the ratio of immune cell densities

We next investigated variability in the ratios of the densities of different cell phenotypes which can predict the nature of the immune response. Specifically, we considered the ratio of CD8α cell density to FoxP3 cell density (CD8α:FoxP3), and the ratio of GZMB cell density to FoxP3 cell density (GZMB:FoxP3), which are typically calculated to assess the extent of anti-tumor versus immunoregulatory T cell abundance in the tumor [20–24]. Figures 4A and 4B show heatmaps for CD8α:FoxP3 and GZMB:FoxP3 ratios, respectively, for all step sections across all animals in different tumor models. For display purposes, the ratios for each tumor model are normalized to one so that results across different tumor models can be presented in the same heatmap. For instance, for the EMT6 model, the CD8α:FoxP3 ratio is relatively constant at different step sections within individual tumors and among tumors from different animals, which correlates with the relatively low intra- and inter-tumor variabilities of 17.7% and35.9% observed, respectively, for CD8α:FoxP3 ratio in this model (Figure 4C). In contrast, for the KPC-GEM tumor model, considerable variability was observed in the CD8α:FoxP3 ratio within individual tumors and among tumors from different animals (e.g., see animal 1, 3 and 5), correlating with the relatively high intra- and intertumor variabilities of 41.7% and 73.1% observed, respectively, in this model (Figure 4C). Similar correlations can be drawn from the analysis of intra-tumor and inter-tumor variability of GZMB:FoxP3 ratios in the different tumor models. Specifically, intra-tumor variability for the GZMB:FoxP3 ratio is relatively low (CV~20%) for the CT26 and EMT6 tumor models and is marginally higher in the other tumor models. Further, inter-tumor variability for GZMB:FoxP3 ratio for the KPC-GEM and KPC-orthotopic tumor models are consistently higher than those for CT26 and EMT6 tumor models.

**Figure 4.**
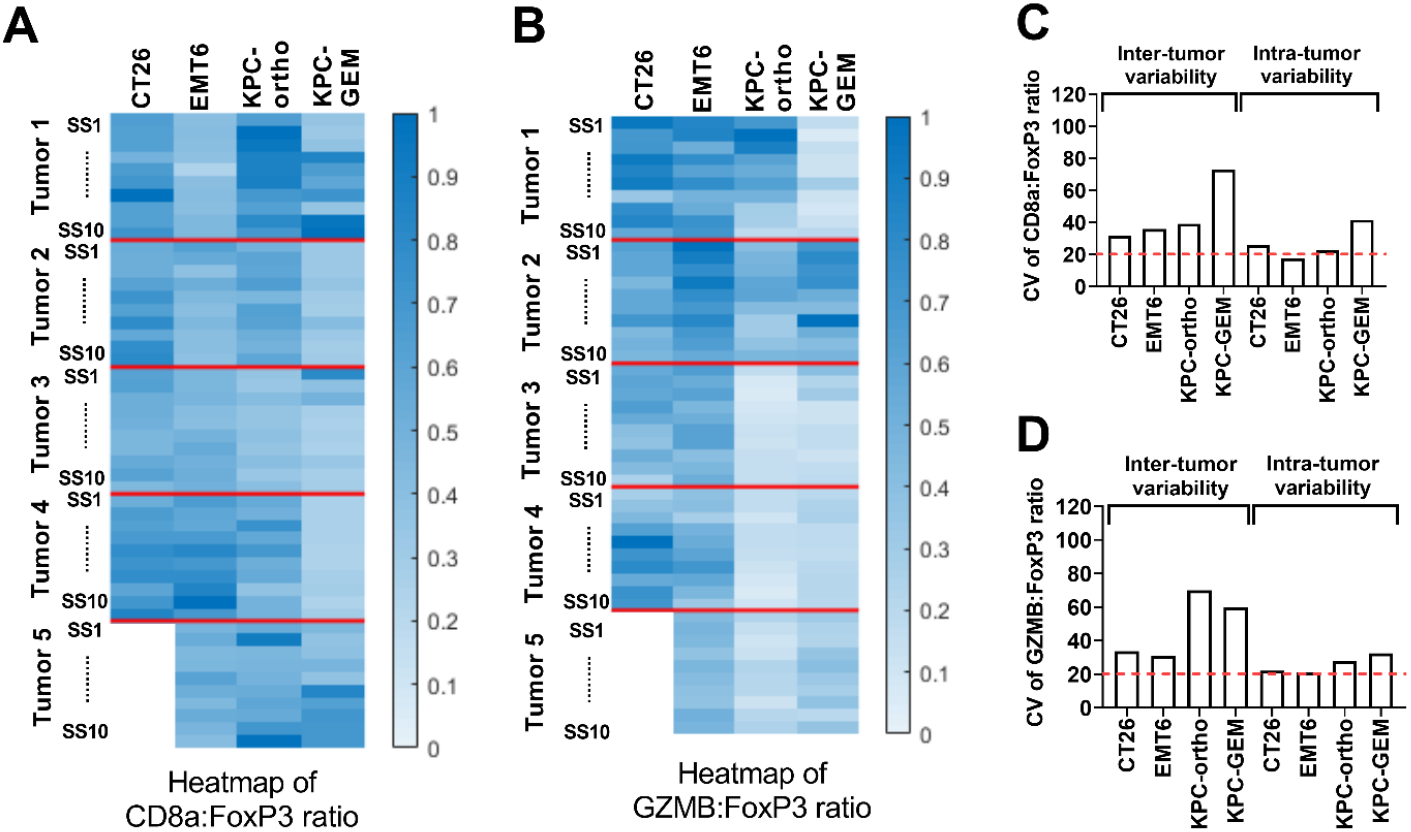
Intra-tumor and inter-tumor variability of CD8α:FoxP3 ratio and GZMB:FoxP3 ratio in syngeneic murine tumor models. Panels A and B show heatmaps depicting the ratios of immune cell densities for CD8α:FoxP3 and GZMB:FoxP3, respectively, in different tumor models across all step sections and animals. The data in the heatmap is normalized in an analogous manner to the heatmap shown in Figure 2A. Panels C and D show the quantification of intra-tumor and inter-tumor variability for CD8α:FoxP3 ratio and GZMB:FoxP3 ratio, respectively, in different tumor models. The calculation of intra-tumor and inter-tumor variability for the ratio of immune cell densities is identical to that of the calculation for immune cell density which was described in Figure 2.

It should be noted that the inter-tumor variability of the immune cell ratios is relatively low when compared to the inter-tumor variability of the corresponding immune cell densities (Figure 2). This is especially the case for CT26 and EMT6 tumor models, where inter-tumor variabilities of CD8α:FoxP3 and GZMB:FoxP3 ratios are ≤35%, whereas the inter-tumor variabilities of CD8α, FoxP3 and GZMB cell densities are ≥60%.

### Pairwise correlation of immune cell biomarkers in different tumor models

We next investigated the pairwise correlation of cell density for all biomarkers in different tumor models. Specifically, for a given pair of biomarkers we calculated the Spearman’s correlation coefficient between the cell density estimates obtained from all step sections among all tumors within a given model. For the CT26 and EMT6 tumor models, we observed relatively high pairwise correlations between T cell-associated biomarkers CD3, CD4, CD8α, FoxP3 and GZMB (Table 1). In contrast, in KPC-orthoptic and KPC-GEM tumor models, T cell-associated biomarkers show poor to moderate correlations (Table 1). For instance, the Spearman’s correlation between CD8α and FoxP3 cell densities in CT26 and EMT6 tumor models are 0.88 and 0.84, respectively, whereas for the KPC-orthotopic and KPC-GEM tumor models, the Spearman’s correlation between CD8α and FoxP3 cell densities are, 0.47 and −0.24, respectively. This suggests that in the CT26 and EMT6 tumor models there is a compensatory mechanism wherein an increase in CD8α cell density results in a proportional increase in FoxP3 cell density. Moreover, this partly explains the relatively low inter-tumor variability that we observed for the CD8α:FoxP3 ratio in these two tumor models and not in the KPC-GEM tumor model, despite the relatively high inter-tumor variability of the CD8α and FoxP3 cell densities that was observed in all the models we evaluated. In contrast to T cell-associated markers, the Spearman’s correlation between myeloid cell markers is less distinct in all of the tumor models. Specifically, pairwise correlations among CD11b, F4/80 and GR1 is either weakly positive or weakly negative except for the EMT6 tumor model, for which the Spearman correlation between GR1 and CD11b is 0.66.

Similar results were also observed for the other DIA endpoint IHC area% (Supplementary Table 1). More specifically, for any given pair of biomarkers, the Spearman’s correlation coefficient between cell densities and the Spearman’s correlation coefficient between IHC area% values were consistently close to each other in all tumor models considered here (supplementary Figure 8).

### Relative proportion of immune cell biomarkers within and across tumor samples

The relative proportion of different immune cell populations in the tumor tissue is typically used to characterize the tumor microenvironment as hot, cold, immunosuppressive, T cell rich, myeloid-cell rich, etc., [6, 7]. As these characterizations are based on bulk measurements of immune cell infiltrates, we next investigated whether the relative proportion of the nine immune cell biomarkers at different depths within and among tumor samples remains the same. We took advantage of our tissue sampling strategy (Figure 1B), where the ordering of the IHC antigens was preserved in the serial sections cut at each 100-micron step. This allowed us to calculate the correlation between two sets of immune cell density estimates *CD3_k,i_, CD4_k,i_*, *CD8_k,i_* …, *GZMB_k,i_* and *CD3_m,j_*, *CD4_m,j_*, *CD8_m,j_* …, *GZMB_m,j_* obtained from animals k and m at steps i and j, respectively, within a given tumor model. Here, *CD3_k,i_* denotes the CD3 cell density from the k^th^ animal at the i^th^ step, *CD4_k,i_* denotes the CD4 cell density from the k^th^ animal at the i^th^ step, and so forth. Figure 5 illustrates the results of this analysis, depicted in heatmaps of the Pearson’s correlation coefficients for different tumor models. For all tumor models, the correlation coefficient is relatively high (i.e., ρ varies between 0.8 and 1) in the diagonal blocks of the heatmaps, which pertain to the relative proportion of immune cells from step sections that are within a given tumor. In the case of the CT26 (Figure 5A) and EMT6 (Figure 5B) tumor models, we also see relatively high correlation coefficient in the off-diagonal blocks of the heatmaps, which pertain to the relative proportion of immune cells from step sections that are within a given tumor. This suggests that the relative proportion of the biomarkers is well maintained within and among tumors in CT26 and EMT6 tumor models. In contrast, for the KPC-orthotopic (Figure 5C) and KPC-GEM (Figure 5D) tumor models, the correlation coefficient is lower (ρ varies from 0.6 to 0.8) in the off-diagonal blocks of the heatmaps. This suggests that there is greater variation in the relative proportion of immune cells in the KPC-orthotopic and KPC-GEM tumor models among tumors when compared to CT26 and EMT6 tumor models.

**Figure 5.**
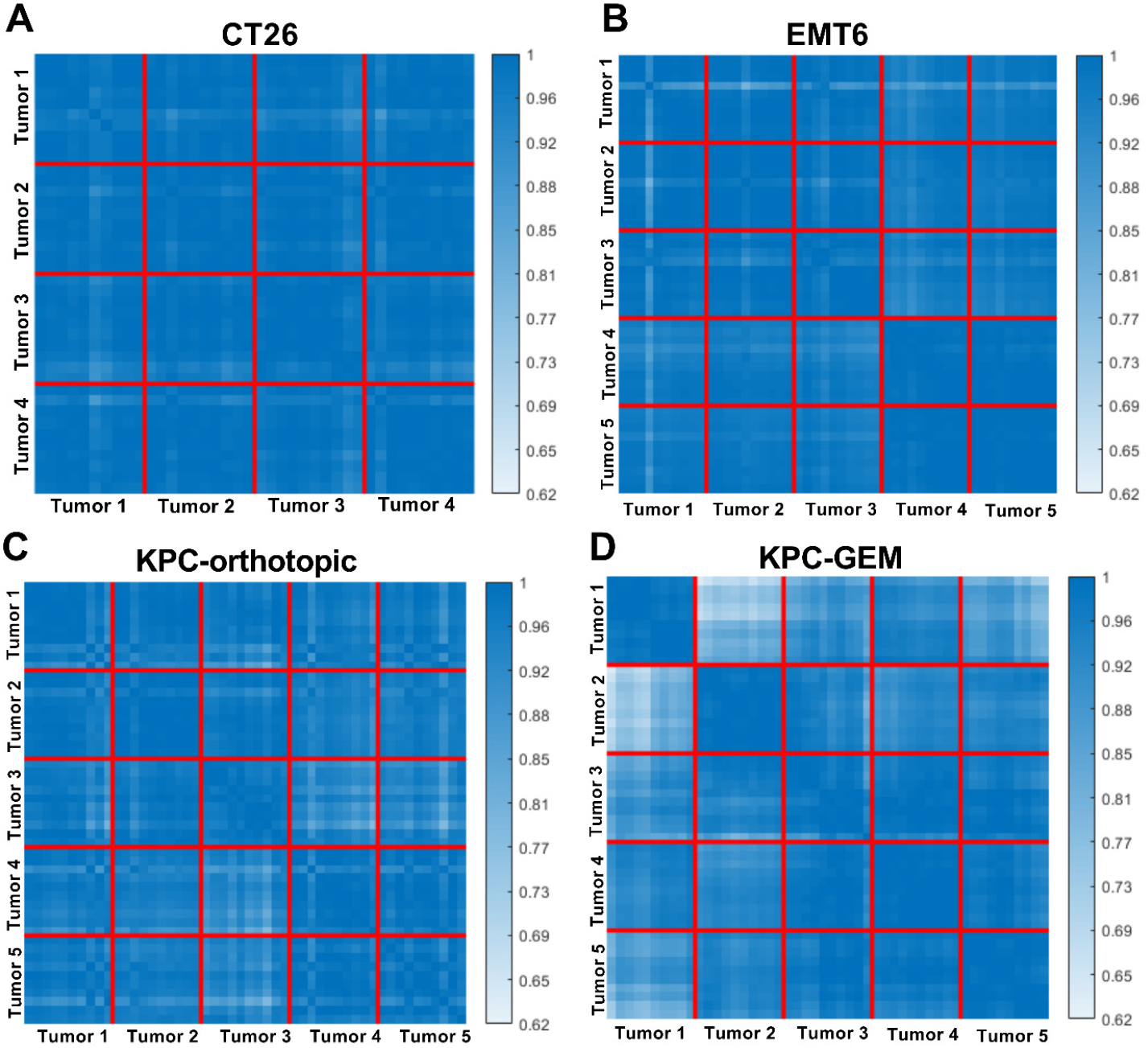
Relative proportions of immune cells within and across tumors. Panel A shows the CT26 tumor model heatmap for the pairwise Pearson correlation coefficient of immune cell density estimates for all nine biomarkers between two step sections that are within and among tumors for a given tumor model. Panels B, C and D show the same for EMT6, KPC-orthotopic and KPC-GEM tumor models, respectively. For each model, the colormap is globally scaled so that the heatmap color coding can be compared within and across animals.

## DISCUSSION

Murine syngeneic tumors are indispensable model systems in preclinical immuno-oncology research. While immunologic heterogeneity has long been recognized as an important factor that can influence treatment outcome, there is a severe paucity of data concerning the nature of this heterogeneity in murine tumor models. Our present study provides a quantitative characterization of the variability in the density and proportion of immune cell infiltrates within and among tumor samples. Specifically, by using multiple tumor models that vary in levels of complexity and translational relevance, our results address longstanding questions concerning immunologic heterogeneity in murine tumors.

Our observation that inter-tumor variability was the dominant source of variation in immune cell density in the tumor models we evaluated suggests animal-to-animal variability is a major contributor, likely arising due to the stochastic nature of tumor evolution which in turn impacts immune cell abundance. The relatively low intra-tumor variability that we observed for most of the biomarkers considered here suggests that a single tumor section is typically adequate to quantify immune cell abundance. The robust agreement between the CV values from the retrospective analysis of IHC-DIA data and the tumor samples analyzed in this study suggest that the small group size (n = 4 or 5 tumors/model) was adequate to recapitulate the inter-tumor variability that was observed from a large aggregate of data pooled from several studies.

The strong correlation that we observed between the two DIA endpoints(i.e., cell density and IHC area%) for all the biomarkers suggest that either of these endpoints can be used to assess immune cell abundance in murine tumors. This is also consistent with the analogous observations made for the inter-tumor and intra-tumor variability of the two DIA endpoints. From a technical standpoint, the computation of cell density requires the detection of individual cells which can sometimes be challenging, for example, due to nuclear overlap, incomplete membrane labeling, the presence of combined membrane and cytoplasmic localization, etc. In such cases, the use of IHC area % could be beneficial as this DIA endpoint is a pixel-based metric that does not rely on nuclear segmentation and/or cell boundary delineation.

Our power analysis revealed that the minimum group size to detect a certain fold-change in cell density can vary depending on the biomarker in question and the tumor model employed. This then raises a practical question as to how one should select the optimal group size for a study that seeks to reliably detect a fold-change in cell density estimates for multiple biomarkers. Ideally, one could select the largest group size to detect a certain fold change in cell density. However, in many situations this may not be feasible; perhaps taking the median of the minimum group size for the different biomarkers could be an acceptable tradeoff in these instances.

CT26 and EMT6 are relatively fast growing, cell line-derived tumors that are typically implanted subcutaneously. In contrast, KPC-GEM tumors are spontaneously occurring tumors of the pancreas that mimic different stages of evolution of human pancreatic cancer. The KPC-orthotopic model used is a cell line-derived tumor from the KPC-GEM tumor model which was implanted orthotopically. Our observation that CT26 and EMT6 tumors but not KPC-orthotopic and KPC-GEM tumors had consistently low intertumor variability for CD8α:FoxP3 ratio and GZMB:FoxP3 ratio raises an important question as to whether this difference is driven by tumor extrinsic (e.g., site of implantation) or tumor intrinsic (e.g., immunogenicity, regulatory mechanisms, etc.) factors.

It has been shown that in several murine tumor models the patterns of T-cell infiltration are comparable between subcutaneous and orthotopic tumors [25, 26], although these and other studies have reported differential responses to immunotherapy depending upon the site of implantation [25–28]. Indeed, the site of implantation could influence the tumor microenvironment and, in turn, the response to immunotherapeutic agents. For instance, the presence of a dense stroma with abundant extracellular matrix deposition is a characteristic feature of pancreatic tumors [29] including the KPC-orthotopic and KPC-GEM tumor models used here [13]. Not surprisingly, collagen, which is the most abundant extracellular matrix protein, is known to affect T cell infiltration and to selectively regulate expression of T cell related genes, including *Foxp3* and *Gzmb* [30]. This may dictate the compensatory increase in FoxP3+ cells with increasing CD8α+ cells in the pancreatic tumor models which we observed in the CT26 and EMT6 tumors. Taken together, these observations suggest that both tumor-intrinsic and tumor-extrinsic factors are likely contributing to the observed differences in the immune cell ratios in the various tumor models considered here.

The insights gained from the analysis of the relative proportion of immune cells in different tumor models is distinct from those inferred from the assessment of inter-tumor variability. Specifically, the calculation of inter-tumor variability is based on individual immune cell densities for a given biomarker across all step sections and all tumors for a given model. In contrast, quantification of the relative proportion of immune cells takes into consideration the immune cell densities of all biomarkers between a pair of step sections within the same tumor or among different tumors in a given tumor model.

Our observation that the relative proportion of immune cells is well preserved among tumor specimens in CT26 and EMT6 models but not in KPC-orthotopic and KPC-GEM models highlights the effects of the tumor microenvironment and the site of implantation on patterns of immune cell distribution and the anti-tumor immune response. An implication of this result is that immune cell composition is generally not preserved among tumor specimens in the KPC-orthotopic and KPC-GEM tumor models. Importantly, this highlights the existence of compositional heterogeneity of immune cells which is distinct from the animal-to-animal variability that already exists in these models due to variations in the immune cell density of individual biomarkers among different tumors. This may also partly explain the generally poor response that has been reported for immunotherapeutic agents in orthotopic and GEM tumor models relative to subcutaneous allografts [9, 25–28].

A shortcoming of our analysis is that our tissue sampling depth was constrained to 1 mm due to the amount of tumor tissue that was available for step sectioning. This was driven by several practical considerations such as the requirement to trim the tissue before embedding in paraffin (e.g. to remove skin and host connective tissue) and the preference to use smaller tumors to limit the formation of necrotic regions especially at the tumor core which severely affects overall tissue integrity.

In conclusion, our study provides a comprehensive analysis of immunologic heterogeneity in murine tumor models. We anticipate that these results will provide guidelines for designing preclinical studies for drug discovery and development.

## AUTHOR CONTRIBUTIONS

SR conceived and designed the study, SO and TC resourced the study and provided logistical support, JA and DT performed the IHC experiments and slide scanning, NS generated and characterized the KPC cell line, JW performed orthotopic tumor implantation, AO, SM and GB performed image analysis, SR and DL performed data and statistical analysis, and SR wrote the manuscript.

## ACKNOWLEDGMENTS

We thank Lisa Manzuk and David Looper for assistance in animal handling, Renee Huynh for assistance with sectioning of FFPE tumor blocks, and Timothy Affolter for critical reading of the manuscript.

