## Supplementary Figures and Tables for "Characterizing intra-tumor and inter-tumor variability of immune cell infiltrates in murine syngeneic tumors"

### SUPPLEMENTARY MATERIAL

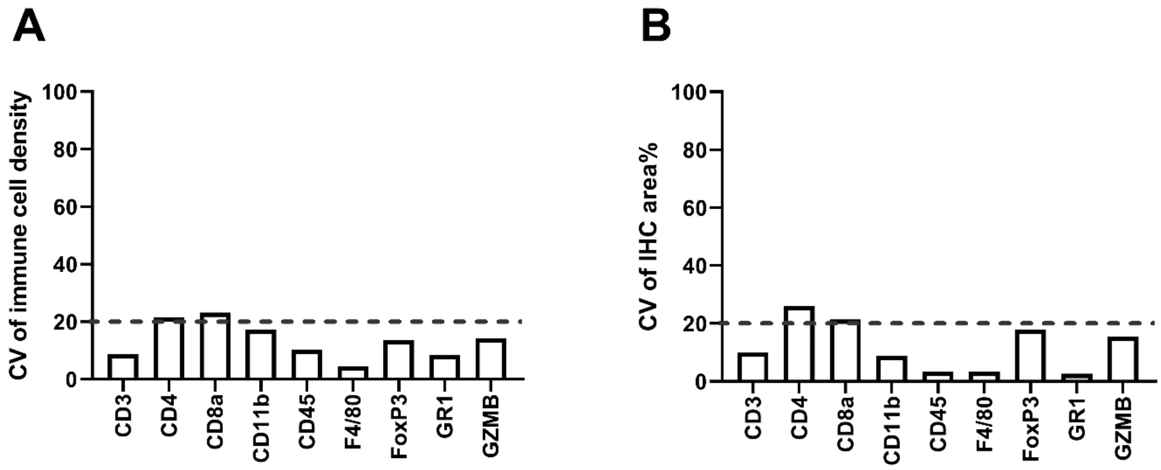


**Supplementary Figure 1**. Results of the precision study for the immune cell biomarkers. Panel A shows the CV of immune cell density for each of the 9 biomarkers, while panel B shows the same for IHC area %. For each biomarker, the CV of a given DIA endpoint was calculated from 15 adjacent serial sections. The dashed line indicates a CV of 20%.

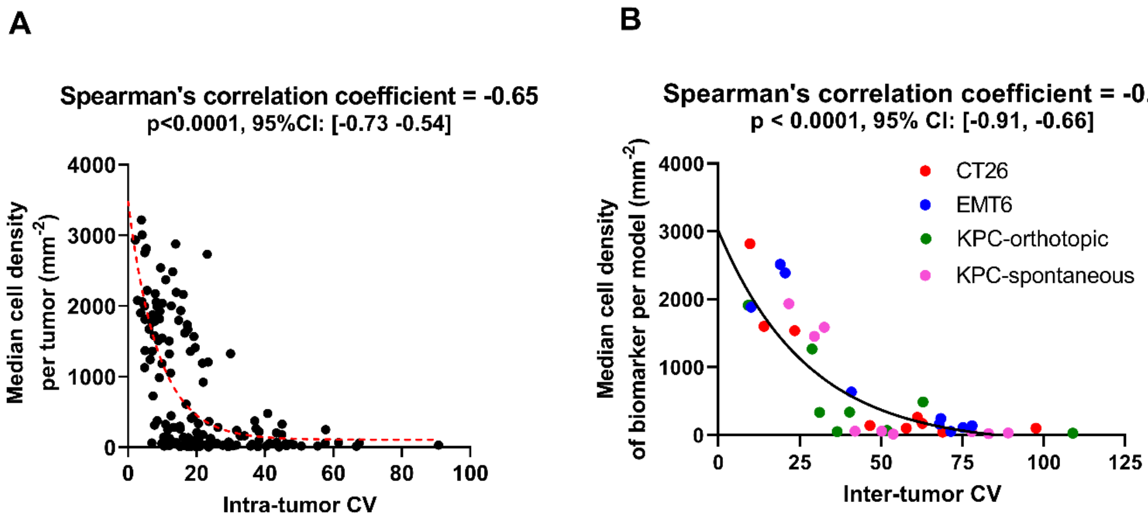


**Supplementary Figure 2**. Correlation between immune cell density and intra/inter-tumor variability. Panel A shows the inverse correlation between median cell density per tumor of every biomarker and its corresponding intra-tumor CV. Panel B shows the inverse correlation between the median cell density of all biomarkers across all step-sections in all tumor specimens versus the corresponding inter-tumor CV, plotted for each tumor model.

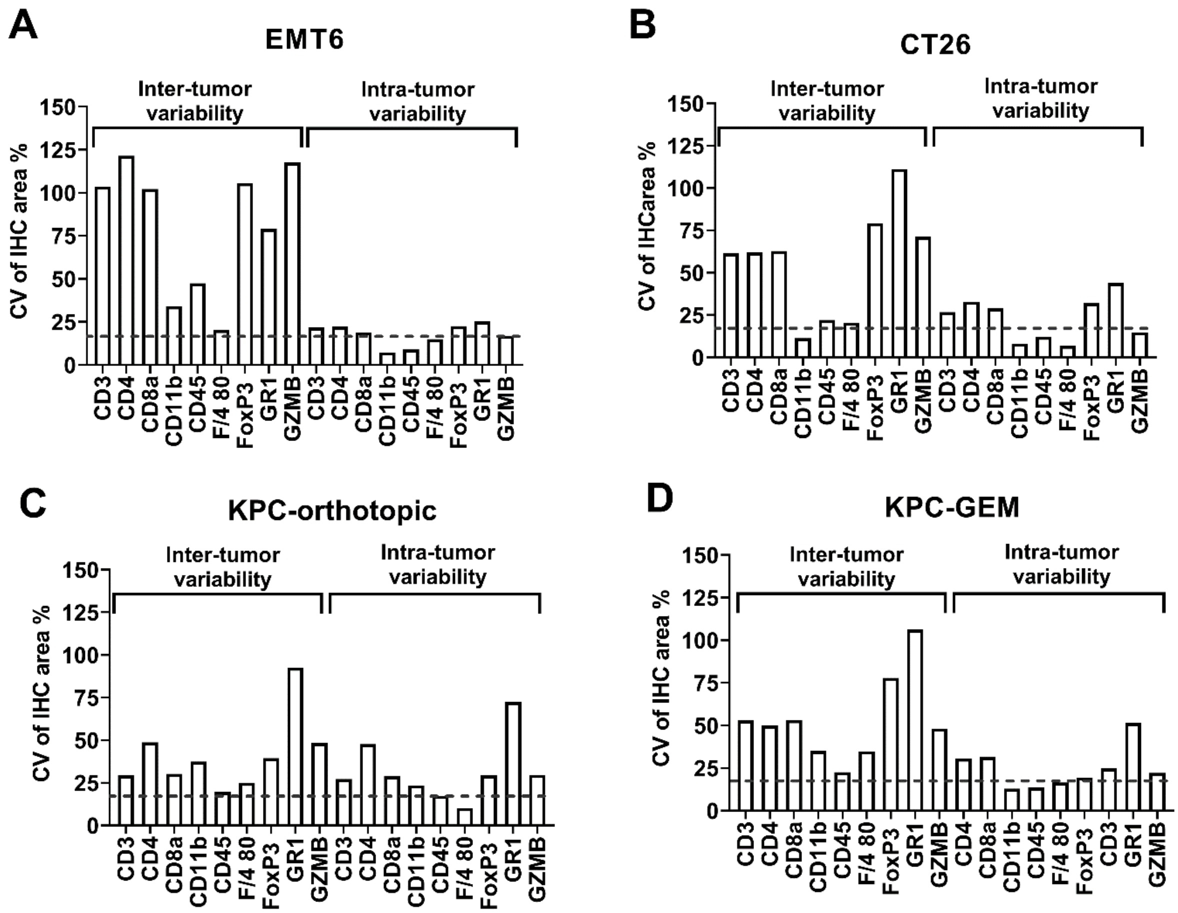


**Supplementary Figure 3.** Inter-tumor and intra-tumor variability in IHC area % for different biomarkers in different tumor models. The dashed line indicates a CV of 20%.

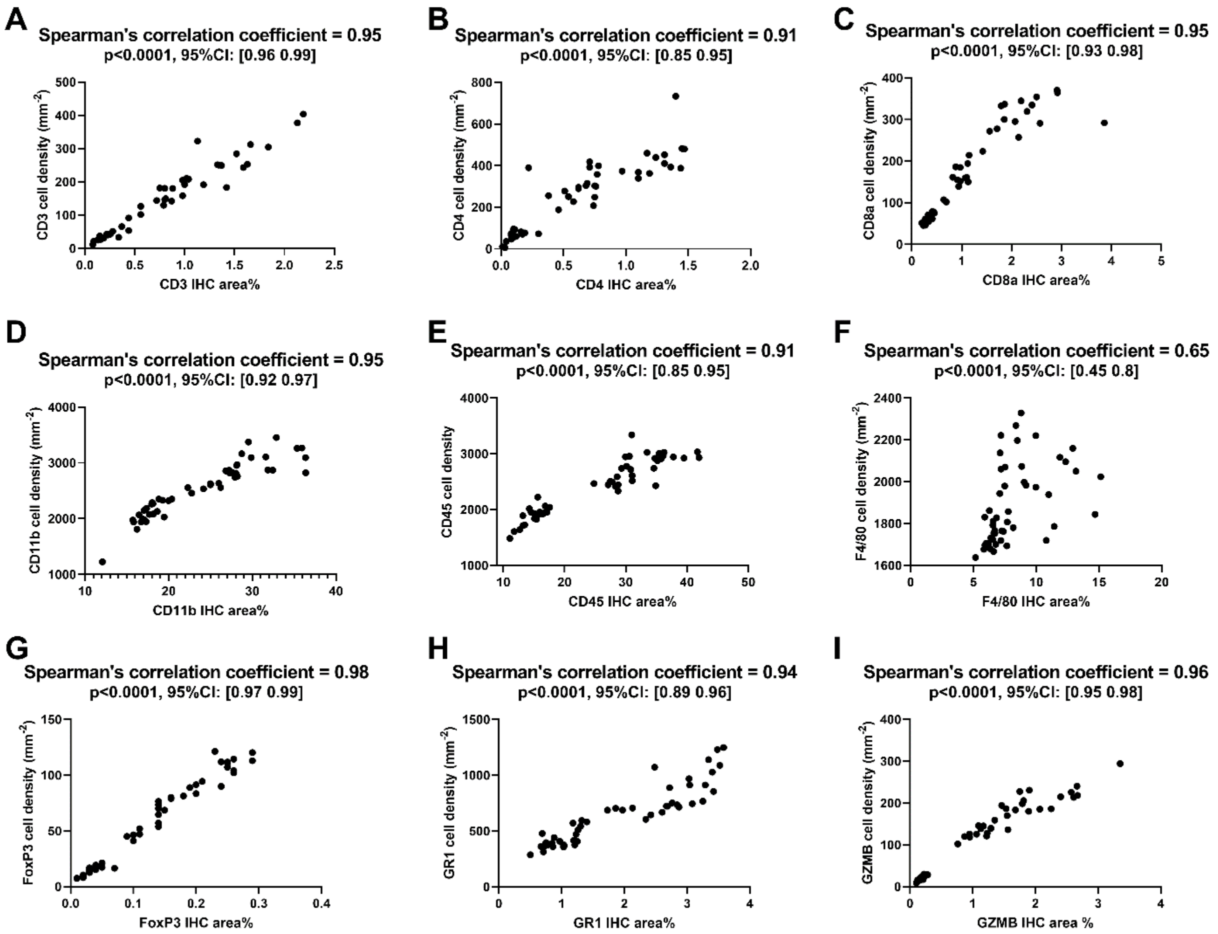


**Supplementary Figure 4.** Plots of IHC area % versus cell density for each biomarker in the EMT6 tumor model. Calculation of Spearman's correlation coefficient between cell density and IHC area% revealed a statistically significant positive correlation for each biomarker (p-value and 95% confidence interval are shown for each biomarker).

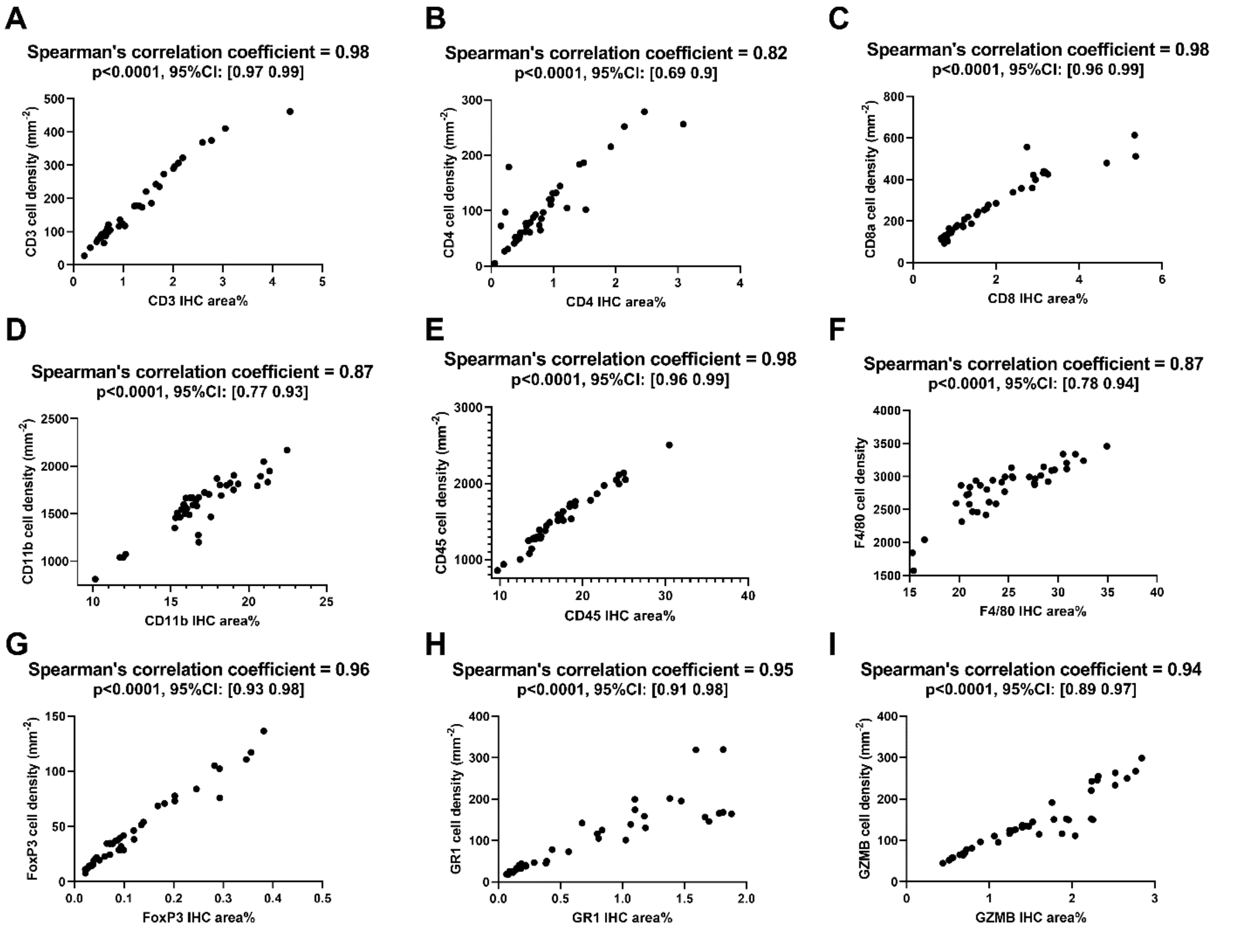


**Supplementary Figure 5**. Plots of IHC area % versus cell density for each biomarker in the CT26 tumor model. Calculation of Spearman's correlation coefficient between cell density and IHC area% revealed a statistically significant positive correlation for each biomarker (p-value and 95% confidence interval are shown for each biomarker).

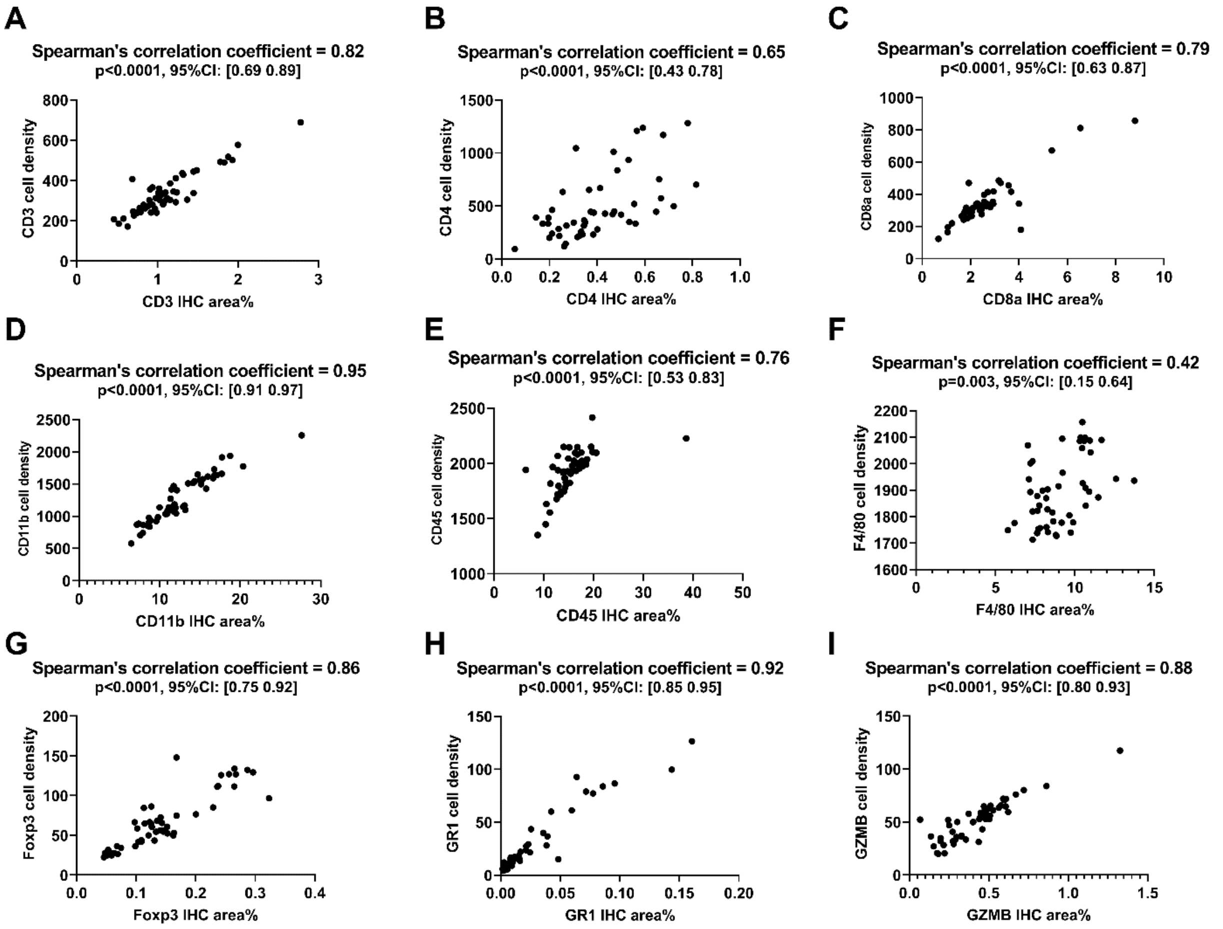


**Supplementary Figure 6**. Plots of IHC area % versus cell density for each biomarker in the KPC-orthotopic tumor model. Calculation of Spearman's correlation coefficient between cell density and IHC area% revealed a statistically significant positive correlation for each biomarker (p-value and 95% confidence interval are shown for each biomarker).

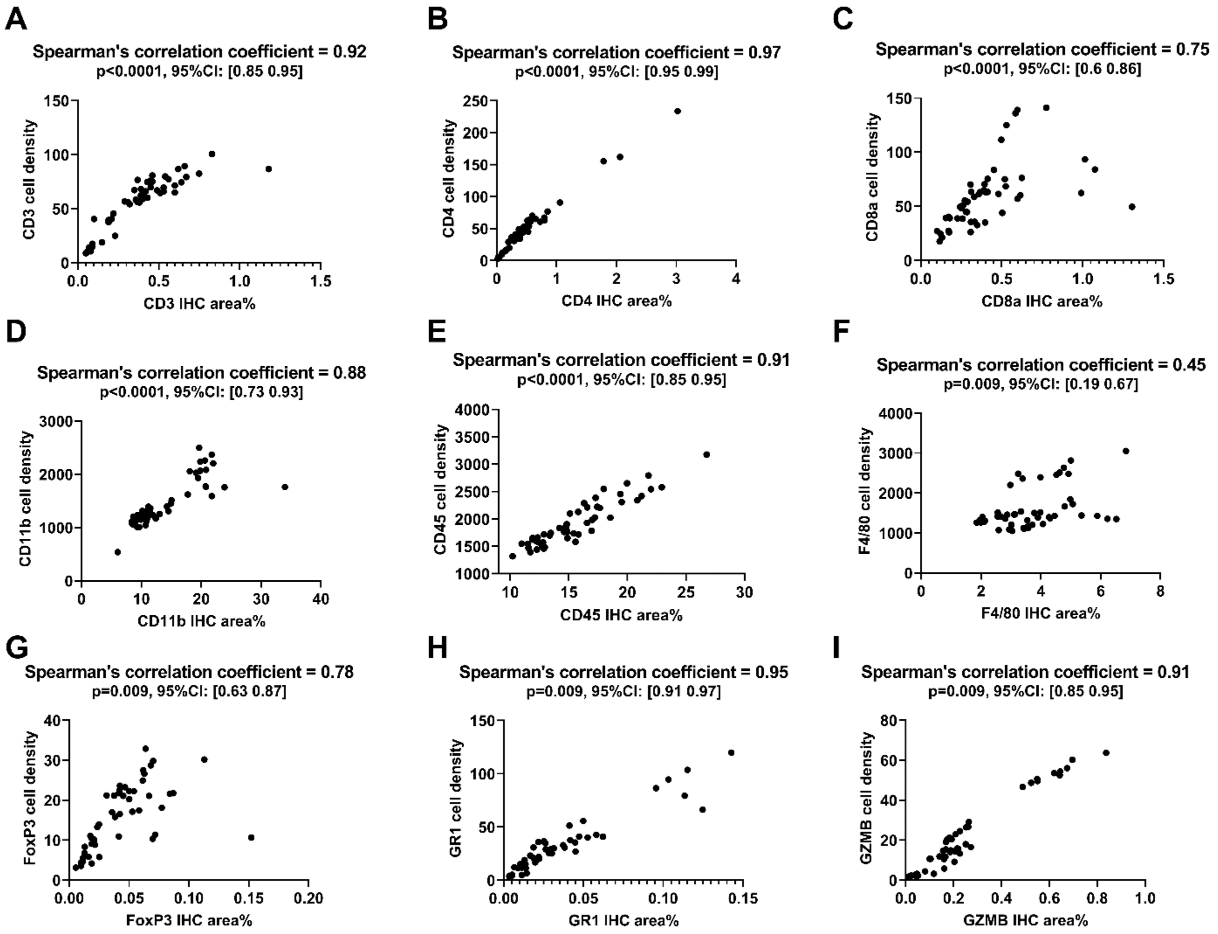


**Supplementary Figure 7**. Plots of IHC area % versus cell density for each biomarker in the KPC-GEM tumor model. Calculation of Spearman's correlation coefficient between cell density and IHC area% revealed a statistically significant positive correlation for each biomarker (p-value and 95% confidence interval are shown for each biomarker).

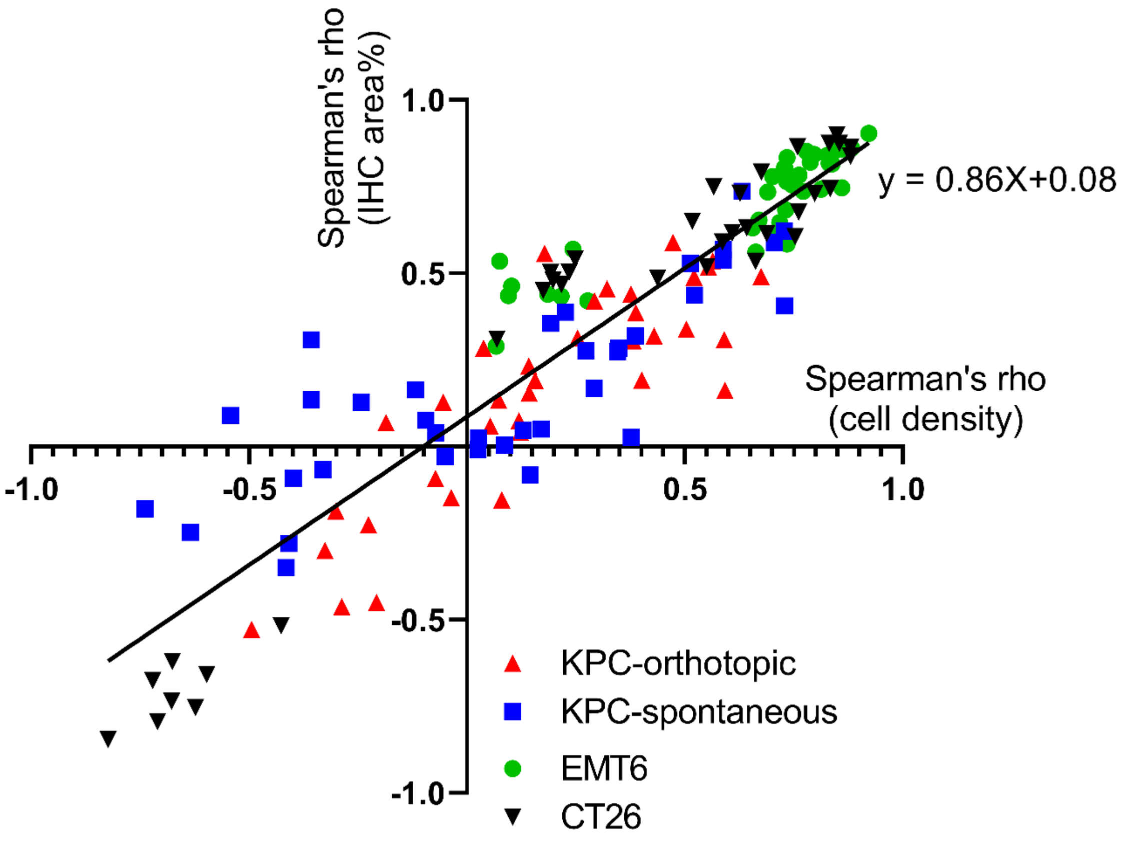


**Supplementary Figure 8**. Plot of the Spearman's correlation coefficient of immune cell density (Table 1) versus the Spearman's correlation coefficient of stain area % (Supplementary Table) for a given biomarker pair in different tumor models. The straight line shows the linear regression analysis between the two Spearman correlation coefficients ($R^2 = 0.84$).

**Supplementary Table 1.** The tables list the pairwise Spearman's correlation coefficient between IHC area % for a pair of biomarkers in different tumor models. The color coding from green to red indicates high positive correlation (+1) to high negative correlation (-1).

### CT26

| | CD3 | CD4 | CD8a | CD11b | CD45 | F4/80 | FoxP3 | GR1 | GZMB |
|---|---|---|---|---|---|---|---|---|---|
| **CD3** | 1.00 | 0.86 | 0.90 | 0.50 | 0.63 | 0.73 | 0.87 | -0.79 | 0.74 |
| **CD4** | | 1.00 | 0.88 | 0.54 | 0.53 | 0.59 | 0.86 | -0.62 | 0.61 |
| **CD8a** | | | 1.00 | 0.50 | 0.65 | 0.75 | 0.84 | -0.75 | 0.68 |
| **CD11b** | | | | 1.00 | 0.45 | 0.48 | 0.47 | -0.52 | 0.31 |
| **CD45** | | | | | 1.00 | 0.79 | 0.52 | -0.73 | 0.62 |
| **F4/80** | | | | | | 1.00 | 0.61 | -0.85 | 0.49 |
| **FoxP3** | | | | | | | 1.00 | -0.68 | 0.73 |
| **GR1** | | | | | | | | 1.00 | -0.66 |
| **GZMB** | | | | | | | | | 1.00 |

##### EMT6

| | CD3 | CD4 | CD8a | CD11b | CD45 | F4/80 | FoxP3 | GR1 | GZMB |
|---|---|---|---|---|---|---|---|---|---|
| **CD3** | 1.00 | 0.83 | 0.84 | 0.78 | 0.82 | 0.43 | 0.86 | 0.74 | 0.75 |
| **CD4** | | 1.00 | 0.83 | 0.81 | 0.82 | 0.42 | 0.85 | 0.62 | 0.76 |
| **CD8a** | | | 1.00 | 0.78 | 0.81 | 0.57 | 0.85 | 0.68 | 0.74 |
| **CD11b** | | | | 1.00 | 0.75 | 0.44 | 0.73 | 0.63 | 0.65 |
| **CD45** | | | | | 1.00 | 0.43 | 0.86 | 0.58 | 0.84 |
| **F4/80** | | | | | | 1.00 | 0.46 | 0.53 | 0.29 |
| **FoxP3** | | | | | | | 1.00 | 0.65 | 0.90 |
| **GR1** | | | | | | | | 1.00 | 0.56 |
| **GZMB** | | | | | | | | | 1.00 |

##### KPC-ortho

| | CD3 | CD4 | CD8a | CD11b | CD45 | F4/80 | FoxP3 | GR1 | GZMB |
|---|---|---|---|---|---|---|---|---|---|
| **CD3** | 1.00 | 0.28 | 0.34 | -0.15 | 0.31 | 0.39 | 0.49 | 0.42 | 0.13 |
| **CD4** | | 1.00 | 0.45 | 0.32 | 0.19 | -0.09 | 0.15 | 0.04 | -0.30 |
| **CD8a** | | | 1.00 | 0.53 | 0.52 | -0.15 | 0.59 | 0.06 | -0.46 |
| **CD11b** | | | | 1.00 | 0.31 | -0.45 | 0.19 | -0.23 | -0.53 |
| **CD45** | | | | | 1.00 | 0.13 | 0.44 | 0.07 | 0.07 |
| **F4/80** | | | | | | 1.00 | 0.16 | 0.49 | 0.56 |
| **FoxP3** | | | | | | | 1.00 | 0.30 | -0.19 |
| **GR1** | | | | | | | | 1.00 | 0.23 |
| **GZMB** | | | | | | | | | 1.00 |

##### KPC-GEM

| | CD3 | CD4 | CD8a | CD11b | CD45 | F4/80 | FoxP3 | GR1 | GZMB |
|---|---|---|---|---|---|---|---|---|---|
| **CD3** | 1.00 | 0.57 | 0.36 | 0.28 | 0.32 | 0.09 | 0.74 | 0.02 | 0.54 |
| **CD4** | | 1.00 | 0.04 | -0.08 | 0.17 | -0.09 | 0.62 | 0.08 | 0.59 |
| **CD8a** | | | 1.00 | 0.28 | 0.53 | -0.07 | 0.13 | 0.39 | -0.01 |
| **CD11b** | | | | 1.00 | 0.27 | 0.31 | 0.00 | -0.35 | 0.03 |
| **CD45** | | | | | 1.00 | -0.18 | 0.05 | -0.03 | 0.44 |
| **F4/80** | | | | | | 1.00 | 0.14 | 0.05 | -0.25 |
| **FoxP3** | | | | | | | 1.00 | 0.16 | 0.41 |
| **GR1** | | | | | | | | 1.00 | -0.28 |
| **GZMB** | | | | | | | | | 1.00 |
